# Spatial attention follows category-based attention during naturalistic visual search: evidence from MEG decoding

**DOI:** 10.1101/390807

**Authors:** Elisa Battistoni, Daniel Kaiser, Clayton Hickey, Marius V. Peelen

## Abstract

In daily life, attention is often directed to high-level object attributes, such as when we look out for cars before crossing a road. Previous work used MEG decoding to investigate the influence of such category-based attention on the time course of object category representations. Attended object categories were more strongly represented than unattended categories from 180 ms after scene onset. In the present study, we used a similar approach to determine when, relative to this category-level modulation, attention is spatially focused on the target. Participants completed two tasks. In the first, they detected cars and people at varying locations in photographs of real-world scenes. In the second, they detected a cross that appeared at salient locations in an array of lines. Multivariate classifiers were trained on data of the artificial salience experiment and tested on data of the naturalistic visual search experiment. Results showed that the location of both target and distracter objects could be accurately decoded shortly after scene onset (50 ms). However, the emergence of spatial attentional selection - reflected in better decoding of target location than distracter location - emerged only later in time (240 ms). Target presence itself (irrespective of location and category) could be decoded from 180 ms after stimulus onset. Combined with earlier work, these results indicate that naturalistic category search operates through an initial spatially-global modulation of category processing that then guides attention to the location of the target.

## 1 Introduction

Top-down attentional selection serves to deal effectively with the large amount of visual information present in everyday environments. It does this by prioritizing processing of goal-relevant stimuli (e.g., cars when crossing a road) and ignoring goal-irrelevant stimuli (e.g., trees when crossing a road). To study this top-down selection mechanisms in the laboratory, many studies have used the visual search paradigm, in which participants are instructed to find simple stimuli, such as oriented bars or colored circles, amongst a variety of distracters (Wolfe and Horowitz, 2004). The use of these artificial displays allows for careful control over variables such as the specific position of targets and distracters and the features that distinguish the target from the distracters. This approach has been fruitfully used in M/EEG studies to reveal the temporal evolution of attentional selection in a variety of visual search paradigms. One of the findings from these studies is that the top-down selection of a peripheral target evokes a lateralized response over occipitotemporal and parietal areas, peaking around 200-300 ms after stimulus onset (Luck and Hillyard, 1994; Eimer, 1996; Hopf et al., 2000; Hickey et al., 2009).

However, visual search in simplified displays differs in many ways from visual search in real life. For example, naturalistic search is typically for a familiar object or object category (e.g., “cars") rather than a visual feature. These target objects appear in scenes that are usually cluttered with a variety of distracters that share many low-level features with the target. Furthermore, the visual properties of target and distracter objects in real-world scenes vary as a function of lighting, perspective, occlusion, and distance.

Despite the apparent complexity of naturalistic search, search in natural scenes is surprisingly efficient (Thorpe et al., 1996; Wolfe et al., 2011). There are several reasons for this efficiency. For example, real-world scenes provide a rich visual context that provides information about likely target features (e.g., objects that are far away appear smaller) and likely target locations (e.g., cars appear on roads). Furthermore, objects in natural scenes are positioned in regular configurations, allowing for the grouping of objects into meaningful chunks (Kaiser et al., 2014). The many differences between artificial and naturalistic visual search highlights the importance of examining the temporal evolution of attentional selection in naturalistic conditions.

A recent magneto-encephalography (MEG) study from our group (Kaiser et al., 2016) took this approach, investigating the time course of object category processing in natural scenes as a function of task relevance. In this study, participants detected either cars or people in a large set of natural scenes. Importantly, the same set of scenes was shown in both tasks, such that objects (cars, people) appeared both as targets and as distracters. Data were analyzed using multivariate pattern analysis (MVPA), decoding the processing of within-scene objects using a classifier trained on data from a separate experiment in which isolated exemplars of cars and people were shown.

Averaged across conditions, the category of the objects present in scenes could be decoded from around 180ms after stimulus onset. Crucially, this early stage of decoding fully depended on the behavioral relevance of the object: early decoding was only possible when the object was the target of the search. These findings show that top-down attention quickly modulates the processing of object category, at around the time that object categories are first being extracted from scenes. However, unlike earlier M/EEG studies investigating attention in simplified displays, the study by Kaiser et al. was not designed to provide information about the spatial component of attentional selection.

In the present study, we closely followed the approach of Kaiser et al. but with significant changes that allowed for measurement of the spatial component of attentional selection. First, targets (cars, people) were presented either in the left or right hemifield so as to elicit lateralized processing. Second, our procedure included an independent experiment in which participants performed a simple detection task in an artificial, non-naturalistic display, reporting the presence of a cross in the left or right hemifield (Fig. 1). We used data from this experiment to train a multivariate classifier to categorize the deployment of spatial attention to the left or right with high temporal resolution. We subsequently used this classifier to detect the deployment of attention to the left or right in data from the main experiment in which participants detected examples of real-world objects in natural scenes.

**Fig.1.**
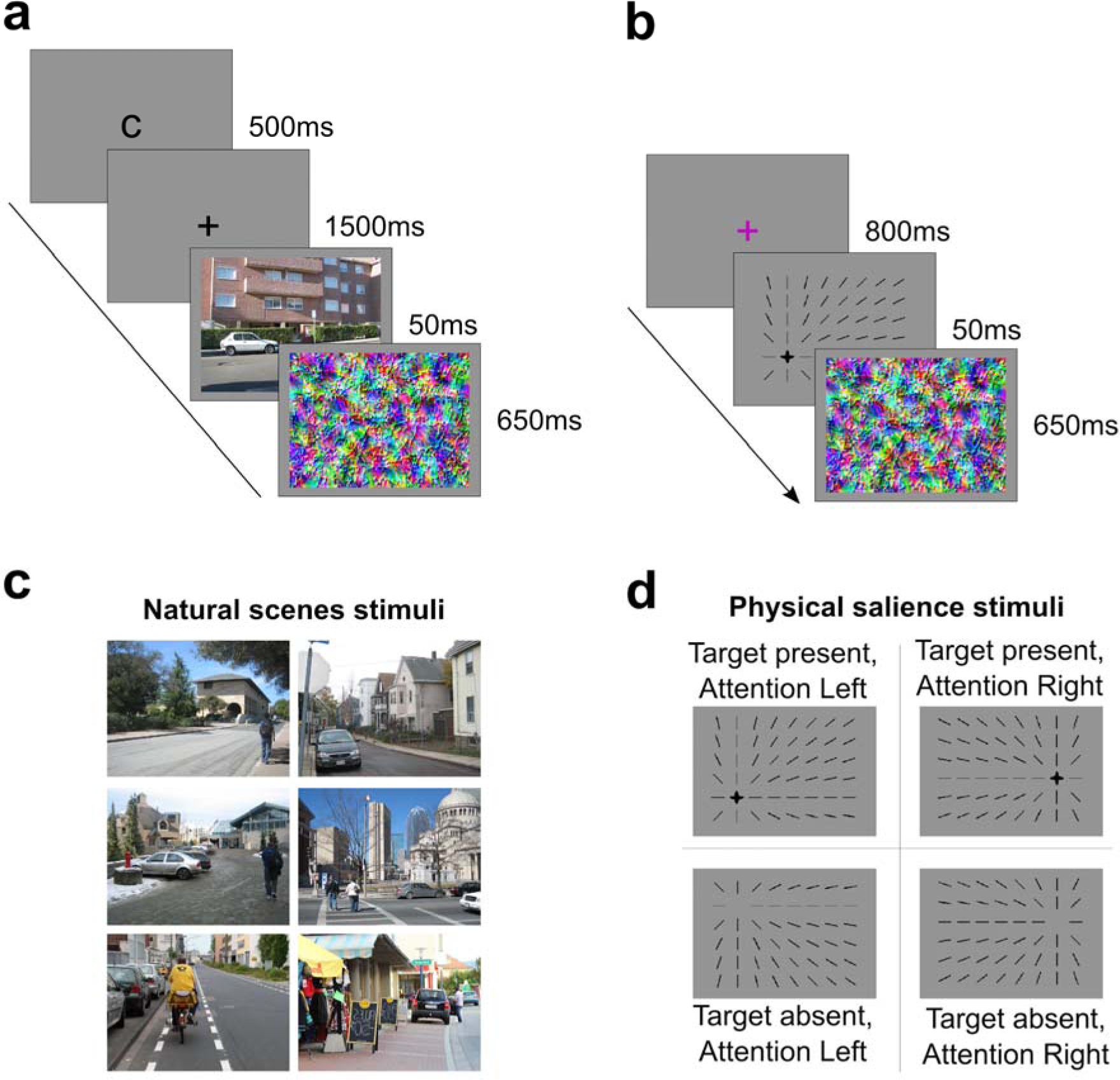
Experimental paradigms. Schematics of paradigms used in (a) naturalistic visual search experiment and (b) physical salience experiment. Example stimuli of (c) the naturalistic visual search experiment and (d) the physical salience experiment.

The objects in the scenes varied as a function of behavioral relevance, allowing us to determine when the location of targets was better decoded than the location of distracters. Importantly, because participants were cued to search for either cars or people in each trial, the same scene stimuli appeared in both target-present and target-absent trials, allowing us to examine neural activity elicited by identical stimuli as a function of whether they currently served as target or nontarget. To exclude the contribution of low-level visual priming, the attended category was symbolically cued, varied on a trial-by-trial basis rather than in blocks, and scenes did not repeat across trials. The cross-decoding approach furthermore allowed us to exclude the contribution of unintended confounds present in natural scenes and thus to relate the deployment of attention to naturalistic stimuli with the deployment of attention to carefully controlled artificial stimuli.

Our findings show that spatial attention is deployed to the target (relative to the distracter) from around 240ms after stimulus onset. Interestingly, information about target presence itself was available from 180ms after stimulus onset, at the same time as the category-based modulation observed by Kaiser et al. (2016). We conclude that spatial attention follows category-based attention during naturalistic visual search.

## 2 Materials and methods

### 2.1 Participants

Data were acquired from 42 healthy participants with normal or corrected-to-normal vision (19 male, mean age M = 26.36 years, SD = 3.75 years). All participants gave informed consent and received monetary compensation. The experiment was conducted in accordance with the Declaration of Helsinki and approved by the Ethical Committee of the University of Trento. Because of a technical problem, no behavioral data was collected for the first three participants.

### 2.2 General experimental procedure

While recording MEG data, participants performed two experiments: a naturalistic visual search experiment in which they detected cars, people or trees in naturalistic scenes (Fig. 1a,c), and a physical salience experiment where they detected the presence of a cross that was made physically salient by converging line elements (Fig. 1b,d). The physical salience experiment was designed to isolate location-specific brain activity patterns, which were used as the training dataset for multivariate classifiers (see below). The full experimental session lasted 80 minutes. Stimuli were back-projected onto a translucent screen located 115cm from the participants. Stimulus presentation was controlled using Matlab 8.0 and the Psychtoolbox (Kleiner et al., 2007).

### 2.3 Naturalistic visual search experiment

In the naturalistic visual search experiment participants reported the presence or absence of a cued target category (cars, people, or trees) in briefly presented photographs of natural scenes by pressing one of two buttons. Participants performed 12 blocks of 48 trials each. The mapping of button to target presence and absence was counterbalanced across participants. As illustrated in Fig. 1a, a letter cue (500ms) displayed at the beginning of every trial indicated the target category (for English-speakers “C” indicated “car”, “P” “person” and “T” “tree"; for Italian-speakers, “M” indicated “macchina”, “P” “persona” and “A” “albero"). After a fixation interval (a “plus” symbol; 1500ms), a natural scene was presented briefly (50ms), and followed by a perceptual mask (650ms). After an additional fixation interval (500ms), participants received feedback (displayed for 500 ms) consisting of 1 or 100 points for correct performance and 0 points for incorrect performance (points were converted to money at the end of the experiment). Trials were separated by a randomly jittered inter-trial interval (rectangular distribution; 1000ms to 2000ms). The average trial duration was 5.2s. The reward feedback manipulation (1 or 100 points) was employed to test a question regarding the effect of reward association on the processing of objects when these appear as distracters. The relevant trials in this context were those where participants searched for trees; these trials are not analyzed or further treated in the current paper. The trials of current interest were those in which participants were cued to detect either cars or people in photographs of real-world scenes that could include one or more exemplars of cars and people (Fig. 1c). Ninety-six scenes contained either cars or people, located on the left or right of the scene. An additional 48 scenes contained both categories (cars and people), where in 24 scenes the two categories appeared on the same side, and in the other 24 scenes they appeared on different sides. In total, the stimulus set consisted of 288 scenes. During the experiment, each scene was presented once in its original version and once flipped horizontally, leading to a total of 576 unique scenes. All pictures were reduced to 480 (vertical) x 640 (horizontal) pixels, subtending 13.5° x 10° of visual angle. Masks of the same size as the scenes (n = 576) were created by superimposing a naturalistic texture to white noise generated at different spatial frequencies, resulting in colored textures. All stimuli were presented centrally and displayed on a grey background.

### 2.4 Phys ca sa ence exper ment

In the physical salience experiment, participants reported the presence or absence of a cross by pressing one of two buttons. The location of the cross was made salient by converging line elements (Fig. 1d) to mimic global contextual cues in natural scenes. Participants performed 2 blocks of 80 trials. The mapping of button to target presence and absence was counterbalanced across participants. Fig. 1b shows the trial structure. After a fixation interval (a pink “plus” symbol presented for 800ms), the line array was presented for 50ms, followed by a perceptual mask (650ms). Trials were separated by a randomly jittered inter-trial interval ranging from 2200ms to 3000ms. The perceptual mask, its timing, and the timing of the stimulus, were identical to those in the naturalistic visual search experiment. Stimuli consisted of 48 black lines on a grey background (displayed on 6 imaginary rows and 8 imaginary columns, each subtending about 1.5° of visual angle), drawn within an area of 13.5° x 10° of visual angle (Fig. 1b). The lines made a position in the display salient; in half of the trials a black cross (the target, of size 1.5° x 1.5°) was presented at this location and in half it was absent. The position of the area within which the target could appear was counterbalanced across 8 possible locations: within the second column (i.e. on the left) or the seventh column (i.e. on the right), the target could appear in one of four positions (in the second, third, fourth or fifth row). All stimuli were displayed on a grey background.

### 2.5 MEG data acquisition and preprocessing

Neuromagnetic activity was recorded using a whole-head MEG system with 102 magnetometers and 204 planar gradiometers (Elekta Neuromag 306 MEG system, Helsinki, Finland). Data were acquired continuously (with online sampling rate of 1000 Hz) and band-pass filtered online between 0.1 and 300 Hz. Offline preprocessing was performed using MATLAB 8.0 and the Fieldtrip toolbox (Oostenveld et al., 2011). Data were epoched from −200 to 500 ms with respect to stimulus onset. No offline filter was applied to the data^1^. Based on visual inspection, and blind to condition, trials and channels containing artifacts (i.e., blinks, eye-movements, or unusually large peak-to-peak amplitudes) were discarded from subsequent analysis. All trials (correct and incorrect) were included in the analysis. Next, data were baseline corrected with respect to the pre-stimulus period (with baseline from −200ms to 0ms) and down-sampled to 100Hz to improve signal-to-noise ratio (Grootswagers et al., 2017). Data from rejected channels were interpolated based on the average of neighboring sensors of the same type.

1 In a previous version of the analysis we applied an offline high-pass filter. This revealed earlier attention effects than without the filter (reported here), possibly reflecting filtering artifacts (Acunzo et al., 2012). These early attention effects did not emerge in subsequent analyses (e.g., reverse cross-decoding, see footnote 2) and were thus deemed unreliable.

2 The cross-decoding analysis was also performed in the reverse direction: training the classifier on the main experiment (separately for target and distracter trials) and testing on the physical salience experiment. This yielded similar results as the analysis reported here: The target-distracter difference on the diagonal emerged after 200 ms, with significant differences from 210ms to 250ms and from 280ms to 380ms. When averaging results of the two cross-decoding analyses, the target-distracter difference was significant (*p*<0.05, corrected for multiple comparisons) from 220ms to 390ms and from 430ms to 500ms.

3 Single sensors did not show reliable attention effects in the current study.

### 2.6 MEG multivariate pattern analysis

All multivariate classification analyses were performed using MATLAB 8.0 and the CoSMoMVPA toolbox (Oosterhof et al., 2016). Single-trial classification was performed separately for every 10ms time bin of the evoked field data of all magnetometers; only data from magnetometers were used as these sensors offered more reliable classification performance than gradiometers in a comparable study (Kaiser et al., 2016). To increase the signal-to-noise ratio, 1000 synthetic trials were created for every condition in both the training and testing sets. Each synthetic trial was created by randomly selecting 5 trials and averaging across these trials. Trials were selected without replacement until the pool of trials was exhausted, such that each trial contributed to a roughly equal number of synthetic trials. Classification accuracy was evaluated by computing the percentage of correct predictions of the classifier. The decoding analysis was repeated for every possible combination of training and testing time, leading to a 50×50 points (i.e. 500 ms x 500 ms with 100Hz resolution) matrix of classification accuracies for every participant. Single-subject accuracy matrices were smoothed using a 3×3 time points averaging box filter (i.e. 30 x 30ms, for the training and testing times, respectively); single-subject accuracy matrix diagonals were smoothed with a 3- point (30 ms) boxcar filter. To determine time periods of significant above-chance classification, a threshold-free cluster enhancement procedure (Smith and Nichols, 2009) was used with default parameters. The multiple-comparisons correction was based on a sign-permutation test with null distributions created from 10,000 bootstrapping iterations and a significance threshold of Z > 1.64 (i.e., p<0.05, one-tailed).

#### 2.6.1 Within-exper ment decoding analyses

A within-experiment decoding procedure was employed to test whether Linear Discriminant Analysis (LDA) classifiers could reliably discriminate MEG activity patterns evoked by stimuli in the left vs. right hemifield. This procedure was performed once within the physical salience experiment and once within the naturalistic visual search experiment. To this end, each of the datasets was divided into two independent subsets of trials, one of which was used as training set and the other as testing set.

#### 2.6.2 Cross-decoding ana yses

In the cross-decoding analysis, LDA classifiers were trained to discriminate between two conditions of interest in the physical salience experiment (MEG patterns evoked by left vs. right stimuli) and employed to discriminate between conditions in the independent naturalistic visual search experiment (MEG patterns evoked by left vs. right objects in natural scenes; Fig. 2a)^2^. This procedure was performed separately for each time point. Classifier testing was performed as a function of the behavioral relevance of objects in scenes, with identical scenes appearing in both target and distracter conditions across participants (Fig. 2b). The difference of the decoding time courses for target and distracter conditions was then tested against zero. It should be noted that the classifier trained on the physical salience experiment can use activity patterns driven by both physical asymmetries and spatial attention shifts. Crucially, however, these can be disentangled in the naturalistic search experiment: in the main comparison, between target and distracter decoding, the same scenes are included as targets and distracters, thus eliminating the contribution of any physical asymmetries.

**Fig. 2.**
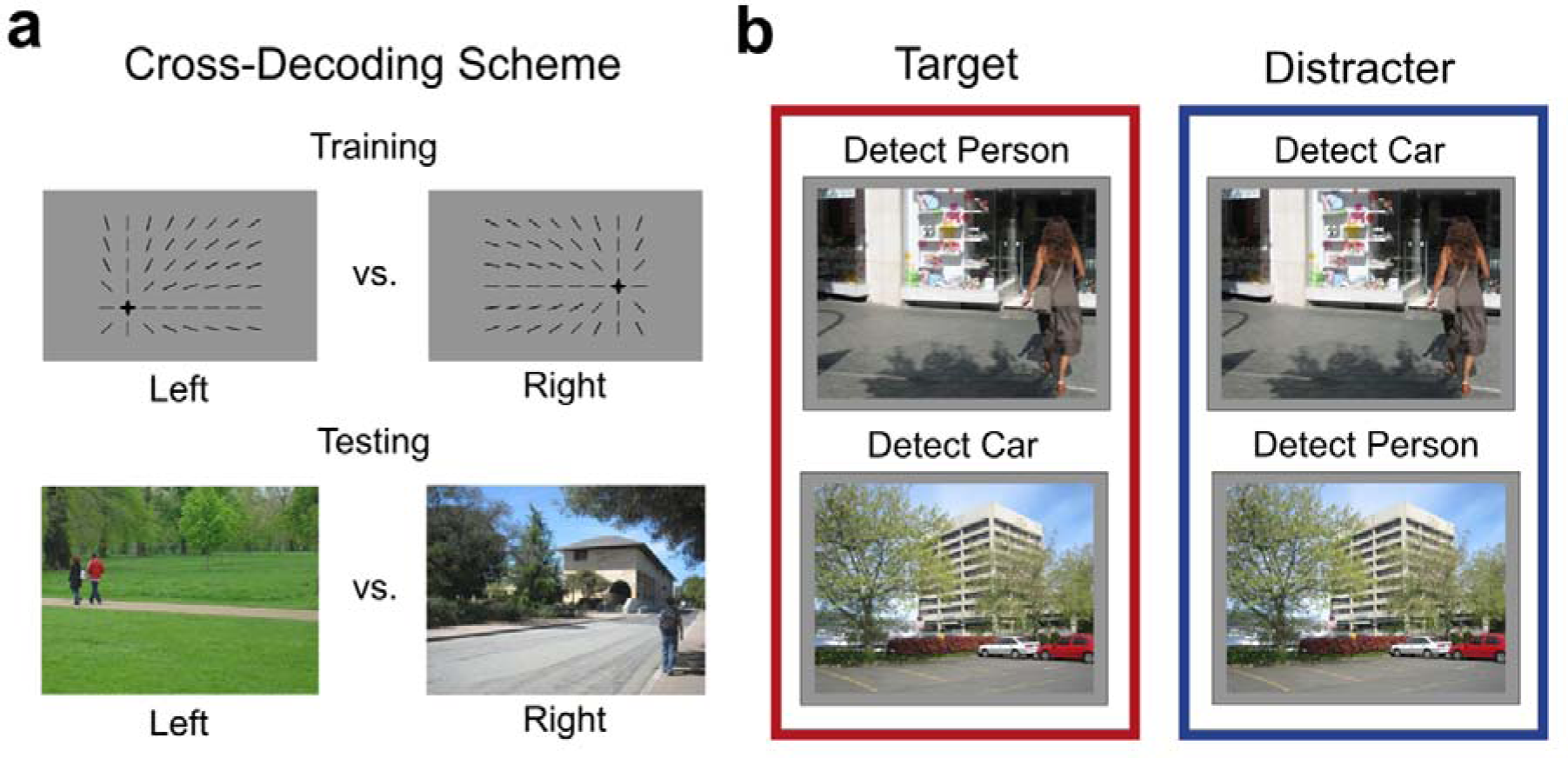
Analysis procedure. Using a cross-decoding approach (a), multivariate classifiers were trained on data from the salience experiment and tested on data from the naturalistic visual search experiment. Classifier testing was performed separately for target and distracter locations (b). Note that the same scenes could appear as target or as distracter, with the only difference being the top-down set of the participant on that trial.

#### 2.6.3 Search lght ana yses

To explore the approximate anatomical location of target and distracter processing, a sensor-space searchlight analysis was performed on consecutive 50ms time windows ranging from 0ms to 500ms post-stimulus. The cross-decoding procedure was performed across the scalp using sensor neighborhoods of 20 sensors each (Kaiser et al., 2016). Each of these neighborhoods was created by defining a neighborhood of 10 adjacent sensors in the left hemisphere that was symmetrically mirrored with corresponding sensors in the right hemisphere, resulting in bilaterally symmetric maps. The searchlight was performed for each 10ms time point, and the results of the individual time points within each 50ms window were averaged to obtain a single searchlight map for that window.

## 3 Results

### 3.1 Behavioral results

Behavioral performance in the naturalistic search experiment showed that the task was sufficiently challenging, with an average response accuracy of around 75% (target present trials: 81%, SD = 10%; target absent trials: 68%, SD = 17%). The average RT was around 500 ms (target present trials: 444ms, SD = 73ms; target absent trials: 555ms, SD = 79ms).

In the physical salience experiment, response accuracy was around 58% (target present trials: 64%, SD = 20%; target absent trials: 52%, SD = 17%). The average RT was around 620 ms (target present trials: 585ms, SD = 190ms; target absent trials: 659ms, SD = 219ms).

### 3.2. Within-experiment decoding results

To determine that MEG activity patterns contained decodable information, and in this way ensure the feasibility of the cross-decoding procedure, in a first analysis we checked whether stimulus location within each experiment was decodable from the data. Within each experiment, multivariate classifiers were trained on a subset of trials to discriminate between left vs. right stimulus location, then tested on a different, independent subset of the data. Significant above-chance decoding was observed within each experiment (Fig. 3). Specifically, decoding within the physical salience experiment was reliable from 100 ms to 500 ms, reaching its maximum accuracy at 330 ms (Fig. 3a). Decoding within the naturalistic visual search experiment was reliable as well, ranging from 90ms to 150ms and from 190ms to 280ms, peaking at 230ms (Fig. 3b). These results highlight that MEG patterns contained information about the stimulus location.

**Figure 3.**
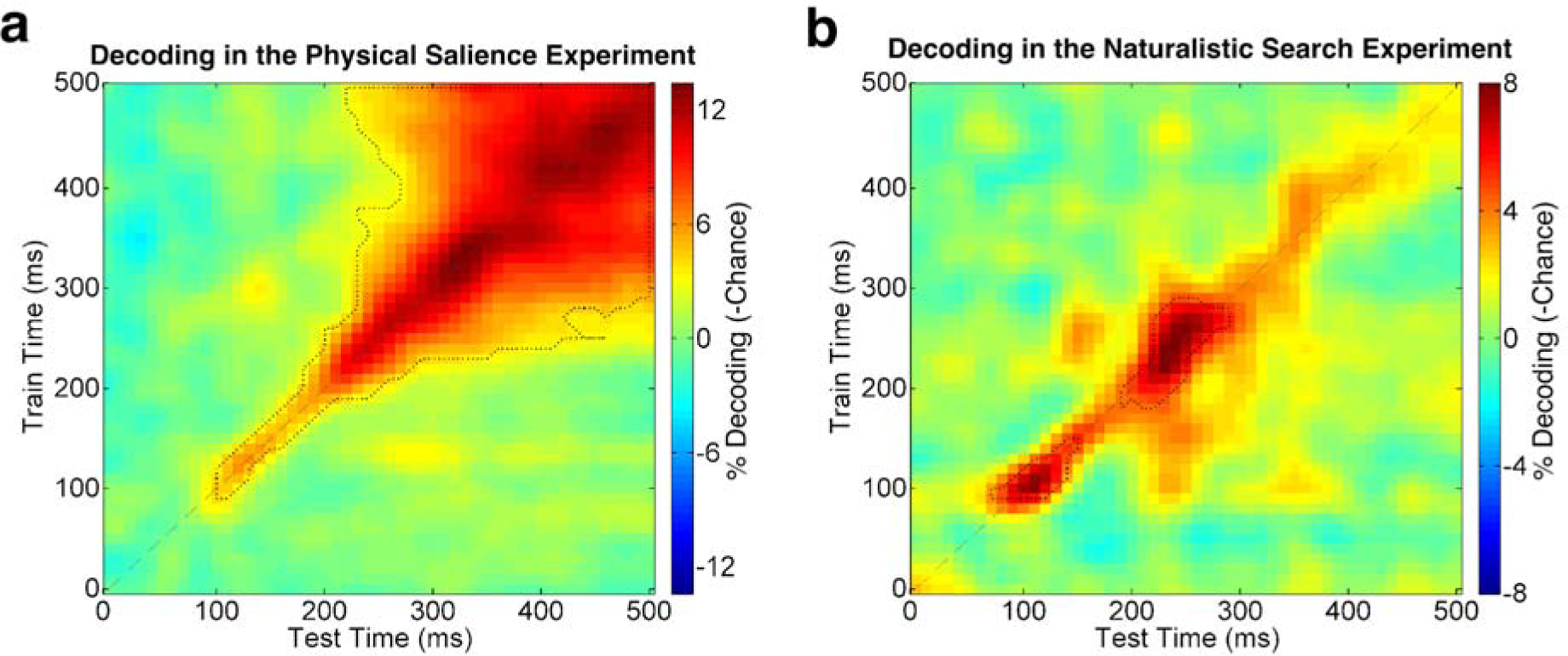
Within-experiment decoding results. Panel (a) shows the time-by-time decoding matrix within the physical salience experiment, panel (b) the resulting matrix within the naturalistic visual search experiment. The outlined areas highlight where decoding accuracy is significantly above chance (p<0.05, corrected for multiple comparisons). Decoding on the diagonal of the matrix in (a) is significantly above chance from 100ms to 500ms, reaching its maximum accuracy at 330ms post-stimulus. Decoding on the diagonal of the matrix in (b) is significantly above chance from 90ms to 150ms and from 190ms to 280ms, peaking at 230ms.

#### 3.3.1 Cross-decoding results

Multivariate classifiers were trained on MEG data from the physical salience experiment and tested on MEG data from the naturalistic search experiment (see Fig. 2). This cross-decoding, averaged across attention conditions (i.e., decoding in target and distracter scenes), was highly reliable from 50ms after stimulus onset, with a first peak at 100ms and a second peak at 260ms (Fig. 4a,b). This result provides evidence for a correspondence between the lateralized processing evoked by the artificial stimuli in the physical salience experiment and the objects in the natural scene experiment. It is worth noting that the decoding peaks of the overall time x time decoding matrix (Fig. 4a) fell on the diagonal, indicating that the temporal evolution of the evoked patterns was similar across the two experiments.

**Figure 4.**
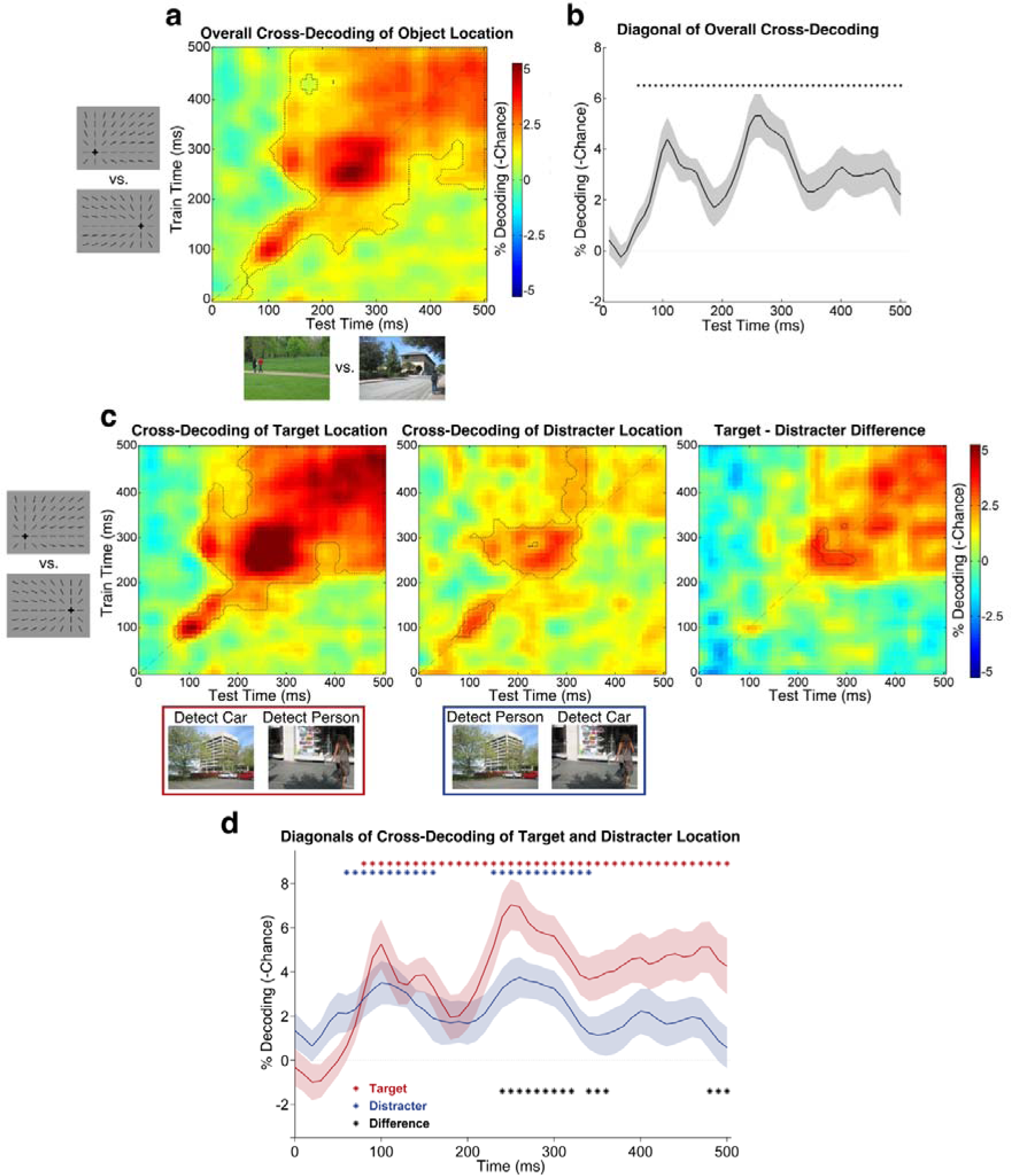
Results of the cross-decoding analysis: time-by-time matrices and time courses of target and distracter conditions. (a) Overall cross-decoding of object location, averaging across target and distracter conditions. The outlined area highlights where decoding accuracy is significantly above chance (p<0.05, corrected for multiple comparisons). Panel (b) shows the diagonal of the overall cross-decoding matrix. This time course is significantly above chance from 50ms after stimulus onset as highlighted by black asterisks (p<0.05, corrected for multiple comparisons). Decoding accuracy was maximum at 100ms and 260ms. Panel (c) shows the time-by-time cross-decoding matrices of the target (left) and the distracter (center) conditions, and their difference (right). Panel (d) shows the time course of decoding target object location (red line) and distracter object location (blue line), reflecting the diagonals of the matrices shown in (c). Shaded coloured areas represent SEM. Target decoding on the diagonal was significantly above chance (p<0.05, corrected for multiple comparisons) from 70ms to 500ms, peaking at 100ms and 250ms (maximum at 250ms). Distracter decoding on the diagonal was significant from 50ms to 150ms, from 220ms to 330ms, peaking at 100ms and 260ms (maximum at 260ms). Target-Distracter difference decoding on the diagonal was significant from 240ms to 320ms, from 340ms to 360ms, and from 480ms to 500ms.

Having established that the location of objects in scenes can be reliably decoded from MEG activity patterns, we next asked when attention modulates this signal. To this end, we separately decoded the position of target objects and distracter objects in otherwise identical scenes. As illustrated in Figure 4c (right panel), decoding of target location was stronger and more reliable than decoding of distracter location. This is clearly illustrated in Figure 4d, which plots the diagonal of these matrices and shows that significant differences emerged from 240ms to 320ms, from 340ms to 360ms, and from 480ms to 500ms.

#### 3.3.2 Searchlight results

To explore the topography of these effects we performed a sensor-space searchlight analysis on consecutive time windows of 50ms each, from 0ms to 500ms post-stimulus. This analysis revealed the time course of the cross-decoding across the scalp, suggesting that the attention effect at 250 ms was primarily driven by lateral posterior sensors before moving more anteriorly (Fig. 5c).

**Figure 5:**
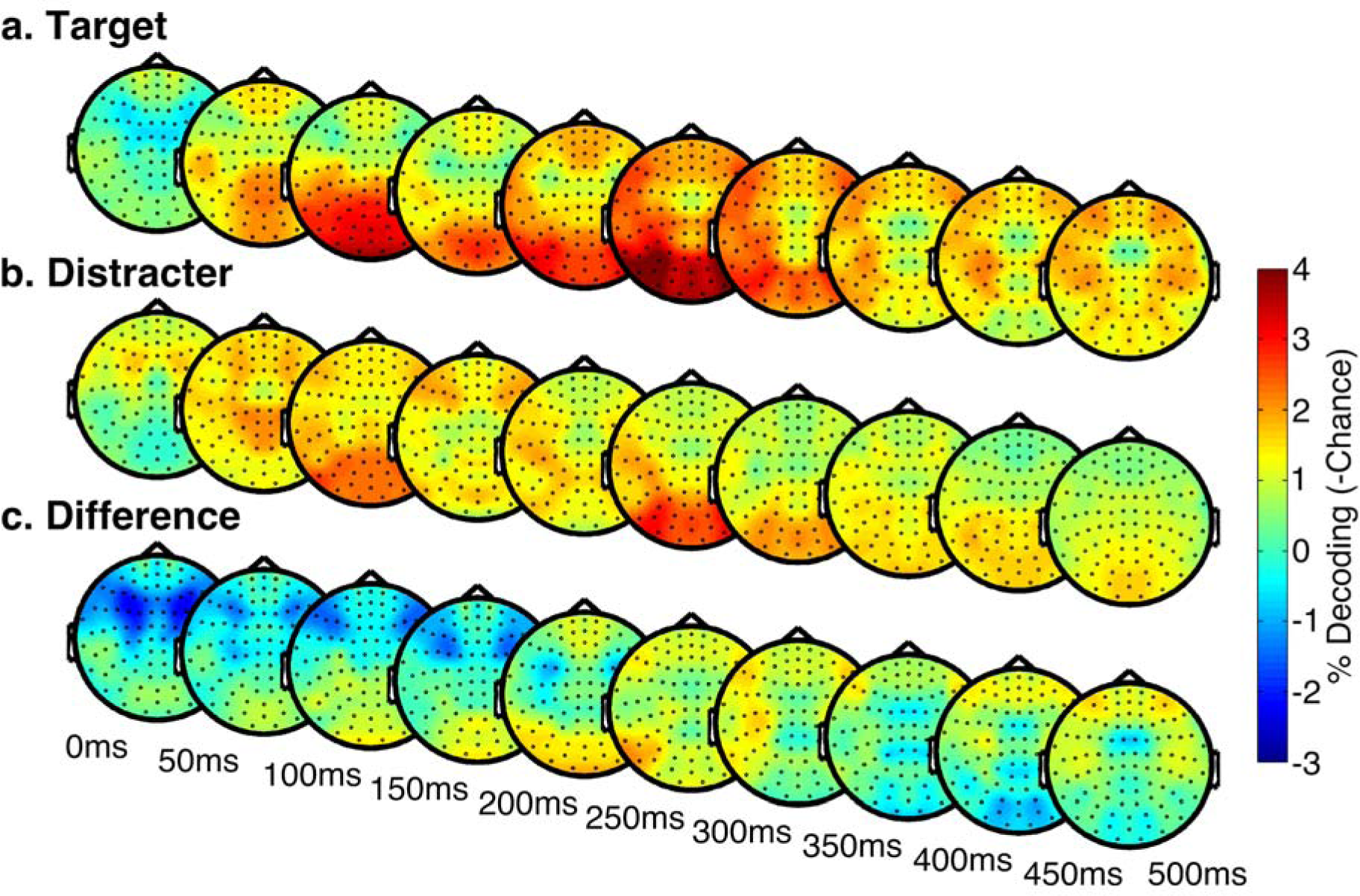
Results of the searchlight analysis. Topographical maps show the results of the cross-decoding searchlight analysis on consecutive time windows of SOms each from Oms to SOOms after stimulus onset, separately for the target condition (a), the distracter condition (b) and their difference (c).

### 3.4. Target-distracter decoding

The cross-decoding analysis provided evidence that spatial attentional selection starts at around 240ms after scene onset, which is later than category-specific attentional modulation in similar tasks, found from 180ms after onset (Kaiser et al., 2016). The presence of category-based attention at 180ms implies that the brain already differentiates target and distracter scenes at that time, thus before the spatial attention effects observed here. To test whether in the current study target and distracter scenes could similarly be differentiated at this time point, we ran an additional analysis within the naturalistic search experiment. In this analysis, we directly decoded the presence of a target (vs. a distracter) in scenes showing either cars or people. Because the only relevant aspect in this analysis was whether the objects were targets or distracters (i.e., matched or mismatched the category of the preceding cue) we averaged across category and location of the objects in the scene. Interestingly, targets could be distinguished from distracters from 180ms after scene onset (Figure 6). The peak was found at 400 ms, shortly before responses were made (mean RT=440 ms in target presence trials). These results indicate that target presence is detected before attention moves to its location.

**Figure 6.**
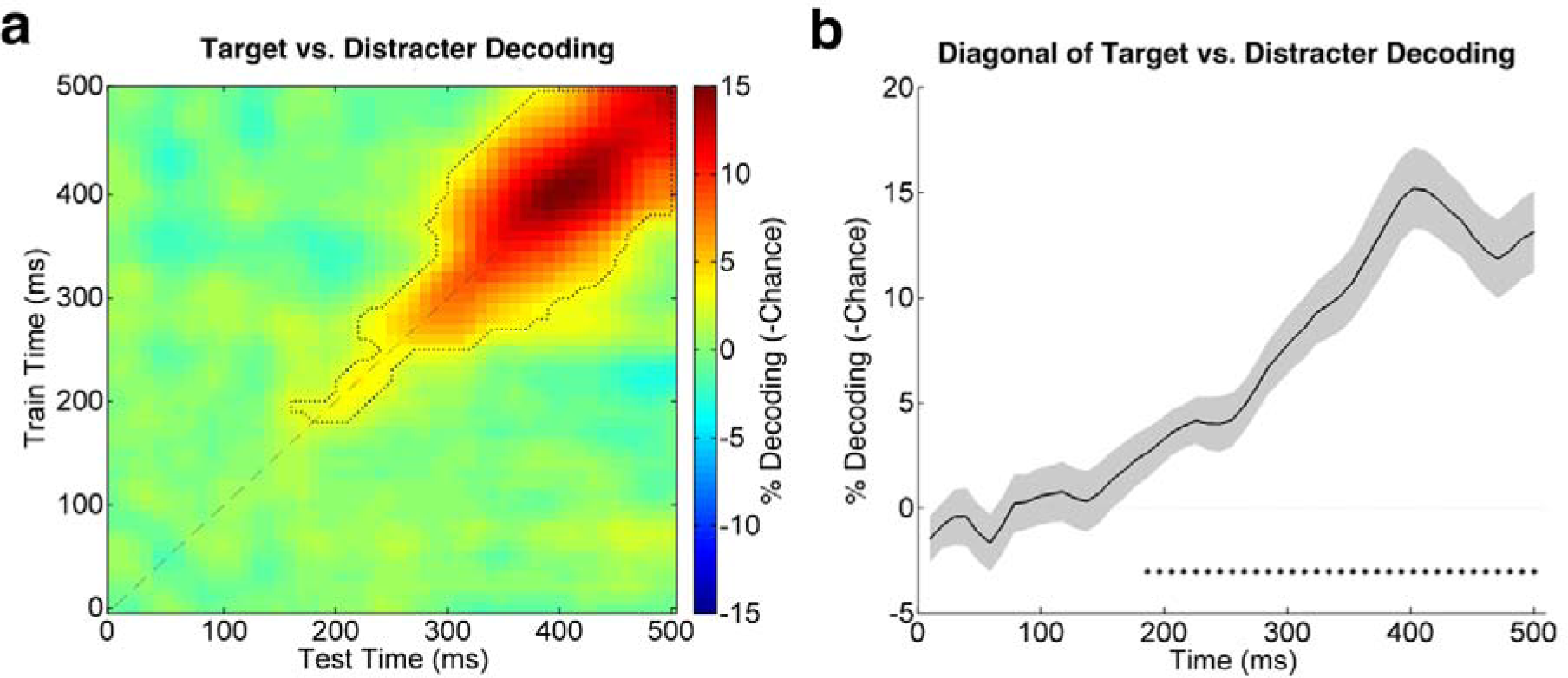
Results of the analysis decoding target vs distracter scenes. (a) Time-by-time matrix of decoding accuracy. The outlined area highlights where decoding accuracy is significantly above chance (p<0 05, corrected for multiple comparisons). Panel (b) shows the diagonal of the decoding matrix. This time course is significantly above chance from 180ms after stimulus onset as highlighted by black asterisks (p<0 05, corrected for multiple comparisons).

## 4 Discussion

The current study investigated the time course of attentional orienting in cluttered natural scenes using multivariate decoding of MEG data. We found that the location of objects in natural scenes can be decoded with high accuracy from MEG activity patterns from 50ms after scene onset. The effect of top-down attention on this decoding arose much later, starting at around 240 ms. Target presence itself (irrespective of location) could be decoded from 180 ms after scene onset. While the decoding of object locations at 50ms clearly reflects a stimulus-driven effect (i.e., presence vs absence of a foreground object), we can be confident that the effects at 180 ms and 240 ms reflect influences of top-down attention: First, the same set of scenes was used for targets and distracters, with target status being determined solely by the match between the scene category and the preceding symbolic cue. Second, we excluded the contribution of bottom-up priming effects because the target category varied unpredictably on a trial-by-trial basis. Taken together with the results of Kaiser et al. (2016), our results indicate that spatial attentional selection follows spatially-global category-based attentional modulation.

The present results are consistent with previous M/EEG studies investigating visual search in artificial arrays. These studies showed that the attentional selection of a target evokes lateralized activity in posterior sensors between 200ms - 300ms after stimulus onset ("N2pc”, e.g., Luck and Hillyard, 1994; Eimer, 1996). Our study indicates that spatial attentional selection in naturalistic search occurs at a similar latency (Fig 4d) and with a similar topography (Fig 5c). This demonstrates an important generalization of previous findings to more naturalistic conditions, despite the differences between artificial and naturalistic search (Wolfe et al., 2011; Peelen and Kastner, 2014) and between univariate and multivariate analysis methods^3^.

The current study complements a recent study that used similar methods to investigate the time course of top-down category-specific attentional modulations in scenes (Kaiser et al., 2016). There, decoding focused on object category processing, with classifiers trained to distinguish exemplars of people and cars and tested on scenes containing exemplars of these categories. Results showed that the category of objects present in scenes could be decoded from around 180ms after stimulus onset. Importantly, this effect was specific to the behaviorally-relevant category from its first emergence, with better decoding of target than distracter category already at 180ms. In other words, top-down attention modulated category-level processing as soon as category information was available.

Our present results show that spatially-specific attention effects - starting at 240ms - emerge after this category-level modulation. This indicates that attention first modulates spatially-global category representations, followed by the spatial selection of the target. This sequence matches that observed in previous studies investigating search for simple features in artificial displays, showing that feature-based attentional modulation precedes spatially-selective enhancement (Hopf et al., 2004; Eimer, 2014). Our results thus support the idea that content-based guidance is not limited to low-level features but can be implemented at higher levels of the visual system as well (Wyble et al., 2013; Hickey et al., 2015; Battistoni et al., 2017; Wyble et al., preprint).

The spatial modulation observed here provides a neural correlate of behavioral findings of attentional capture by objects matching a top-down category-based attentional set (Reeder and Peelen, 2013; Reeder et al., 2015). In these studies, participants searched for cars and people in natural scenes. On a subset of trials, two irrelevant stimuli appeared instead of the scenes. One of these stimuli was quickly followed by a dot that participants were instructed to detect. Results showed that participants were faster to detect the dot when it appeared at the location of a stimulus that shared mid-level features with the target category (e.g., a wheel of a car, or an arm attached to a torso), providing evidence for attentional capture. Importantly, the effect was also observed when the mid-level features appeared at locations that were never relevant to the search task. These findings demonstrate that category-based attention is spatially global and that it guides spatial attention to template-matching stimuli. In conjunction with Kaiser et al. (2016), the current results provide novel insight into the temporal evolution of both these effects.

Interestingly, although spatial attention is captured by template-matching objects, the detection of familiar object categories in natural scenes may not require spatial attentional selection. For example, target-specific EEG responses in these tasks have been observed before 200 ms (Thorpe et al., 1996), more likely corresponding to the category-based modulation observed in Kaiser et al. (2016) than the spatial selection observed here. Similarly, in the current study response patterns evoked by target and distracter scenes differed from around 180 ms after stimulus onset, indicating that target features are detected before spatial attention moves to the target location. Behavioral studies have shown that participants may not be able to localize object categories in natural scenes that have nonetheless been detected (Evans and Treisman, 2005). Others have argued that the detection of familiar object categories may not even require spatial attention at all (Li et al., 2002; see also Stein and Peelen, 2017). These findings suggest that spatially-global category-based attention may be sufficient for detecting target-diagnostic features.

In daily life, however, the detection of category-diagnostic features is often not sufficient for guiding our behavior. Many situations require us to bind features to identify objects at finer levels. For example, we might need to distinguish our red car from our friend's green car, or to find our friend among other people. These tasks require spatial attention to bind features, as elegantly shown by work in neurological patients with parietal damage (Cohen and Rafal, 1991; Friedman-Hill et al., 1995). Thus, while not directly required in the current task, spatial selection may be an integral and obligatory aspect of top-down attention, even when directed to high-level categories (Wyble et al., 2013; Reeder et al., 2015).

To conclude, the current study shows that spatial attentional selection of target objects in natural scenes occurs at around 240 ms after scene onset. This spatial modulation follows an earlier spatially-global categorical attention modulation that provides information about target presence from around 180 ms. Our results are in line with theories of visual search proposing that spatial attention is guided by feature-based selection (Treisman and Sato, 1990; Wolfe, 1994), and importantly generalize this idea to naturalistic search for familiar object categories in natural scenes.

## References

Acunzo DJ, MacKenzie G, van Rossum MCW (2012) Systematic biases in early ERP and ERF components as a result of high-pass filtering. J Neurosci Meth 209:212-218.

Battistoni E, Stein T, Peelen MV (2017) Preparatory attention in visual cortex. Ann N Y Acad Sci 1396:92-107.

Cohen A, Rafal R (1991) Attention and Feature Integration: Illusory Conjunctions in a Patient With a Parietal Lobe Lesion. Psychol Sci 2:106-110.

Eimer M (1996) The N2pc component as an indicator of attentional selectivity. Electroencephalogr Clin Neurophysiol 99:225-234.

Eimer M (2014) The neural basis of attentional control in visual search. Trends Cogn Sci 18:526-535.

Evans KK, Treisman A (2005) Perception of objects in natural scenes: Is it really attention free? J Exp Psychol Human 31:1476-1492.

Friedman-Hill SR, Robertson LC, Treisman A (1995) Parietal contributions to visual feature binding: evidence from a patient with bilateral lesions. Science 269:853-855.

Grootswagers T, Wardle SG, Carlson TA (2017) Decoding Dynamic Brain Patterns from Evoked Responses: A Tutorial on Multivariate Pattern Analysis Applied to Time Series Neuroimaging Data. J Cognitive Neurosci 29:677-697.

Hickey C, Di Lollo V, McDonald JJ (2009) Electrophysiological indices of target and distractor processing in visual search. J Cogn Neurosci 21:760-775.

Hickey C, Kaiser D, Peelen MV (2015) Reward guides attention to object categories in real-world scenes. J Exp Psychol Gen 144:264-273.

Hopf JM, Boelmans K, Schoenfeld MA, Luck SJ, Heinze HJ (2004) Attention to features precedes attention to locations in visual search: evidence from electromagnetic brain responses in humans. J Neurosci 24:1822-1832.

Hopf JM, Luck SJ, Girelli M, Hagner T, Mangun GR, Scheich H, Heinze HJ (2000) Neural sources of focused attention in visual search. Cereb Cortex 10:1233-1241.

Kaiser D, Stein T, Peelen MV (2014) Object grouping based on real-world regularities facilitates perception by reducing competitive interactions in visual cortex. P Natl Acad Sci USA 111:11217-11222.

Kaiser D, Oosterhof NN, Peelen MV (2016) The Neural Dynamics of Attentional Selection in Natural Scenes. J Neurosci 36:10522-10528.

Kleiner M, Brainard D, Pelli D (2007) What's new in Psychtoolbox-3? Perception 36:14-14.

Li FF, VanRullen R, Koch C, Perona P (2002) Rapid natural scene categorization in the near absence of attention. Proc Natl Acad Sci U S A 99:9596-9601.

Luck SJ, Hillyard SA (1994) Spatial filtering during visual search: evidence from human electrophysiology. J Exp Psychol Hum Percept Perform 20:1000-1014.

Oostenveld R, Fries P, Maris E, Schoffelen JM (2011) FieldTrip: Open Source Software for Advanced Analysis of MEG, EEG, and Invasive Electrophysiological Data. Comput Intel Neurosc.

Oosterhof NN, Connolly AC, Haxby JV (2016) CoSMoMVPA: Multi-Modal Multivariate Pattern Analysis of Neuroimaging Data in Matlab/GNU Octave. Front Neuroinform 10.

Peelen MV, Kastner S (2014) Attention in the real world: toward understanding its neural basis. Trends Cogn Sci 18:242-250.

Reeder RR, Peelen MV (2013) The contents of the search template for category-level search in natural scenes. J Vis 13:13.

Reeder RR, van Zoest W, Peelen MV (2015) Involuntary attentional capture by task-irrelevant objects that match the search template for category detection in natural scenes. Atten Percept Psychophys 77:1070-1080.

Smith SM, Nichols TE (2009) Threshold-free cluster enhancement: Addressing problems of smoothing, threshold dependence and localisation in cluster inference. Neuroimage 44:83-98.

Stein T, Peelen MV (2017) Object detection in natural scenes: Independent effects of spatial and category-based attention. Atten Percept Psychophys 79:738-752.

Thorpe S, Fize D, Marlot C (1996) Speed of processing in the human visual system. Nature 381:520-522.

Treisman A, Sato S (1990) Conjunction search revisited. J Exp Psychol Hum Percept Perform 16:459-478.

Wolfe JM (1994) Guided Search 2.0 A revised model of visual search. Psychon Bull Rev 1:202-238.

Wolfe JM, Horowitz TS (2004) What attributes guide the deployment of visual attention and how do they do it? Nat Rev Neurosci 5:495-501.

Wolfe JM, Vo ML, Evans KK, Greene MR (2011) Visual search in scenes involves selective and nonselective pathways. Trends Cogn Sci 15:77-84.

Wyble B, Folk C, Potter MC (2013) Contingent attentional capture by conceptually relevant images. J Exp Psychol Hum Percept Perform 39:861-871.

Wyble B, Hess M, Callahan-Flintoft C, Folk C (preprint) Conceptual content in images triggers rapid shifts of covert attention. bioRxiv.

